# Characterization of gene repression by designed transcription activator-like effector dimer proteins

**DOI:** 10.1101/2020.07.14.202762

**Authors:** NA Becker, JP Peters, TL Schwab, WJ Phillips, JP Wallace, KJ Clark, LJ Maher

## Abstract

Gene regulation by control of transcription initiation is a fundamental property of living cells. Much of our understanding of gene repression originated from studies of the *E. coli lac* operon switch, where DNA looping plays an essential role. To validate and generalize principles from *lac* for practical applications, we previously described artificial DNA looping driven by designed Transcription Activator-Like Effector Dimer (TALED) proteins. Because TALE monomers bind the idealized symmetrical *lac* operator sequence in two orientations, our prior studies detected repression due to multiple DNA loops. We now quantitatively characterize gene repression in living *E. coli* by a collection of individual TALED loops with systematic loop length variation. Fitting of a thermodynamic model allows unequivocal demonstration of looping and comparison of the engineered TALED repression system with the natural lac repressor system.

**Statement of Significance:** We are designing and testing in living bacteria artificial DNA looping proteins engineered based on principles learned from studies of the *E. coli* lac repressor. The engineered proteins are based on artificial dimers of Transcription Activator-Like Effector (TALE) proteins that have programmable DNA binding specificities. The current work is the first to create unique DNA repression loops using this approach. Systematic study of repression as a function of loop size, with data fitting to a thermodynamic model, now allows this system to be compared in detail with lac repressor loops, and relevant biophysical parameters to be estimated. This approach has implications for the artificial regulation of gene expression.

## Introduction

Gene regulation by control of transcription initiation is fundamental to living cells. Interactions between proteins and DNA, and between proteins, can drive RNA polymerase recruitment to, or exclusion from, promoter sequences in DNA. Control is typically through accessory and regulatory proteins, often tuned by post-translational modifications (1–3). The repressive characteristics of eukaryotic and archaeal chromatin suggest the generalization that regulation of eukaryotic and archaeal transcription initiation involves promoter activation, while prokaryotic gene regulation is fundamentally repressive (4). However, both activation and repression of transcription initiation are observed in all three kingdoms of life.

Key insights into the control of prokaryotic transcription initiation were originally gained from classic investigations of the *E. coli lac* operon (5) and the left and right promoters of coliphage *λ* (6). *Lac* control illustrates both repression and activation functions of accessory proteins influencing RNA polymerase binding to the *lac* promoter (3,5,7). The *λ* system similarly illustrates negative and positive control, but also was the first system to demonstrate cooperative binding by clusters of λ repressor proteins locally and through DNA looping (6,8,9).

We have been studying repression of transcription initiation in the LacI repressor system (10–16) because of our interest in understanding how the bending and twisting rigidities of the DNA double helix are managed in living cells (17). The classic studies of Müller-Hill (5,18–21) and Record (22–24) were the first to demonstrate that the *lac* switch features auxiliary (distal) operators in addition to the proximal operator that overlaps the promoter. It was shown that repression of transcription initiation is controlled by the effective concentration of lac repressor at the proximal operator. Effective repressor concentration at this operator is increased by simultaneous binding of bidentate lac repressor tetramer at distal operators. This repression enhancement occurs by cooperativity at a distance via DNA looping (25–29). We have previously studied gene control by assembling elements of the *lac* control switch that allow us to deduce probabilities of DNA looping as a function of DNA length using expression of the *lacZ* gene as the readout (10,13). This approach allows sensitive measurement of biophysical details of DNA looping energetics in vivo at base pair (bp) resolution using ensemble experiments.

Our past studies have illuminated fundamental aspects of the LacI repressor DNA looping mechanism, including the interplay of intrinsic operator affinity (controlled by both DNA sequence and the binding of inducer), operator position, DNA bending and twisting flexibilities, and architectural DNA binding proteins that modify the physical properties of DNA (10–15). With this background, we have recently sought to exploit fundamental principles of the *lac* operon in designing an artificial DNA looping system for application in controlling transcription initiation at any promoter in *E. coli* or other organisms. Our premise is that adequate understanding of the natural *lac* system should enable construction of an artificial system mimicking some of its features, but targeted to regulate arbitrary promoters. We described elements of such an artificial control system based on fusions between designed Transcription Activator-Like Effector (TALE) proteins and dimerization domains controllable by small molecules (30). TALE proteins originating in bacterial pathogens of plants (31,32) employ independent base-specific DNA recognition modules to bind the DNA major groove, allowing engineered targeting. This platform allows us to fuse DNA-binding domains with dimerization domains to create artificial DNA looping proteins. Depending on the choice of dimerization domains, dimerization can be constitutive (as in the present study) or made to be dependent on either the presence or absence of small molecules. Our initial study introduced the design of these sequence-specific DNA-binding proteins, confirmed their ability to act as repressors by targeting *lac* operator sequences, and presented preliminary evidence of repression by DNA looping. This evidence came from the observation that repression was enhanced by a distal (upstream) operator, and enhancement depended on the spacing between proximal and distal operators (a signature of DNA looping). Further, repression depended on TALE protein dimerization to form Transcription Activator-Like Effector Dimers (TALEDs). In the initial study, the sequence symmetry of the idealized *lac* operator meant that TALE monomer binding occurred in either of two orientations, such that each operator spacing could support 2 or 4 competing DNA loops that formed simultaneously, depending on the specific operators and TALEDs being studied. While producing clear evidence for looping enhancement of repression, data quantitation and interpretation were complicated by the potential for multiple TALED-mediated DNA loop geometries.

We now extend our previous results in order to characterize in detail promoter repression by a designed TALED. We measure how gene repression depends on the relative orientation of operators, and unequivocally document DNA looping by measuring its length-dependence at bp resolution for cases where only a single loop conformation is possible at each operator spacing. Importantly, these new coherent data for single loops allow meaningful quantitative thermodynamic modeling of engineered TALED-based gene control elements. This modeling was previously impossible. In turn, this new analysis permits a first systematic comparison with corresponding biophysical parameters obtained from our prior studies the natural *lac* system. This analysis sets the stage for implementation of TALED-directed DNA looping for gene control in other prokaryotic and eukaryotic systems and comparison with other designed approaches (33–35).

## Materials and Methods

### DNA looping reporter constructs

Episomal and plasmid DNA looping constructs (Supplemental Table S1 and Supplemental Figs. S1 and S2) were based on plasmid pJ2280 (30). Episomal spacing constructs were created by modifications of pFW11-null as described in supplemental methods (36,37). Plasmid constructs contain the complete *lacZ* coding sequence downstream of the promoter and operator(s).

### TALE-FKBP protein expression

Cloning of genes encoding designed TALEs involved described methods (38). TALE-FKBP protein expression plasmid was created using a modified version of plasmid pJ1035 (promoter of moderate strength) (37). Plasmid pJ1035 contains the bacterial UV5 promoter with complete −10 and −35 box sequences. See Supplemental Methods and Supplemental Fig. S3 for full details.

### *E. coli* β-galactosidase reporter assay

*LacZ* expression (*E*) was measured using a liquid β-galactosidase colorimetric enzyme assay (39) adapted as previously described (30). Assays were performed with a minimum of 3 colonies repeated on each of 2 days for at least 6 data points. Normalized reporter expression (*E*’), with or without TALE-FKBP, allows for comparisons among experiments:

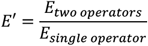

Repression was quantitated in terms of Repression Ratio (*RR*):

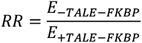

with the contribution to the repression ratio due to free repressor binding at the proximal operator defined as *RR_F_*:

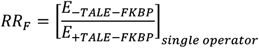

the overall contributions to the repression ratio due to free repressor and DNA looping defined as

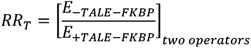

and the contributions to the repression ratio due to DNA looping defined as *RR*’:

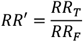

### Data fitting to thermodynamic model

The thermodynamic model of promoter repression used to fit data relating gene expression to the presence and spacing of operator sequences has been previously described (10,13,17). The adaptation of this model to the current analysis is explained in Supplemental Methods. Briefly, the fraction of proximal operator bound by TALED protein as a function of DNA operator-operator length is modeled with six adjustable parameters evaluating the distribution of possible states of the proximal operator through a partition function for the system. Fit parameters give insight into the physical properties of the nucleoprotein loop.

## Results and Discussion

### TALED design

A designed TALED (Fig. 1A, Supplemental Fig. S3) was created to recognize a 15-bp subsequence within the asymmetric *lac* O_2_ operator (38). TALE fusion with a C-terminal FKBP(F36M) mutant domain (Fig. 1A “DD”) in place of the natural TALE transcription activation domain, allows constitutive homodimerization with affinity reported to be 30 μM in vitro (40).

**Fig. 1.**
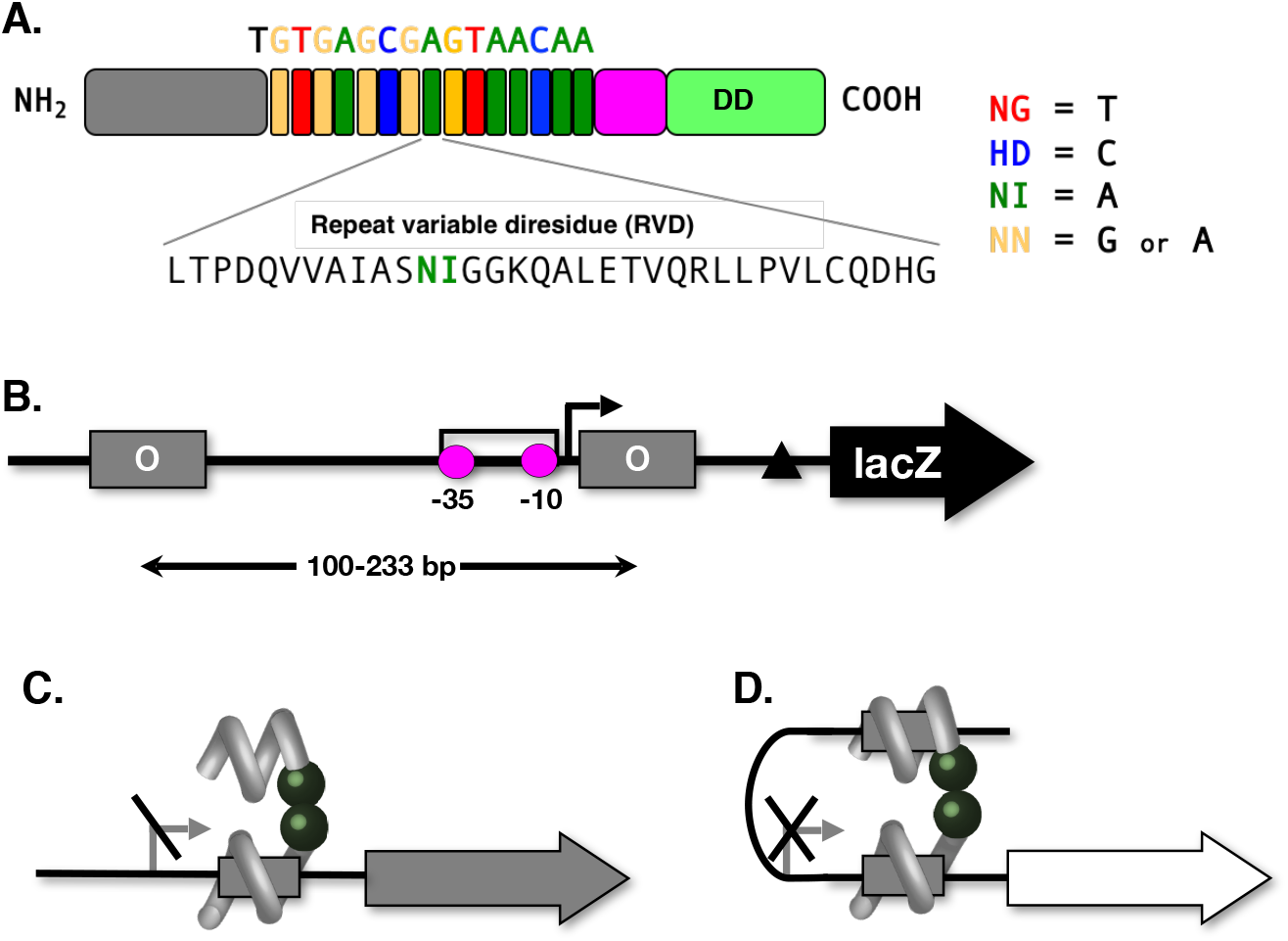
Experimental design. A. TALE protein design and fusion to C-terminal FKBP(F36M) mutant dimerization domain (“DD”). Tandem 34-amino acid repeats (single letter amino acid codes) with programmed base-specific repeat variable diresidue (RVD) domains are indicated in colors. The 15-bp DNA sequence recognized by this TALE protein is indicated above. B. Example of promoter construct design for DNA looping studies. *lac* operators flank a *lac* UV5 promoter (broken arrow shows transcription start site with −10 and −35 sequences indicated) such that the proximal operator is just downstream of the promoter and the distal operator is at various distances (measured operator center-to-center) upstream. The *lacZ* gene acts as reporter and the Shine-Dalgarno sequence is indicated (triangle). C. Schematic of weak repression by TALED binding only the proximal operator in an unlooped configuration. D. Strong repression by TALED-dependent DNA looping for an example operator configuration.

### *lac* looping model systems

As in our prior studies, we demonstrated and characterized engineered DNA looping in vivo using promoter-reporter constructs of the form shown in Fig. 1B. The *lac* UV5 promoter driving *lacZ* is flanked by identical O_2_ operators (Supplemental Fig. S1) derived from the *lac* operon. We intentionally chose to target *lac* operators to allow use of promoter-reporter constructs previously created for analysis of DNA looping by LacI repressor, and to facilitate direct comparison of results. The center-to-center operator spacing is systematically varied to monitor the relationship between the energetically-unfavorable DNA bending and twisting required for TALED-driven looping, and the transcriptional readout. TALED binding to the isolated proximal operator inhibits promoter function (Fig. 1C), and this repression is enhanced by increasing local TALED concentration and promoter distortion by looping (Fig. 1D).

### Effect of operator orientation on TALED-dependent gene repression in vivo

Our prior study (30) involved TALED recognition of a symmetrical O_sym_ *lac* operator, complicating interpretation of results because the directional TALE protein can bind the symmetrical operator in either of two orientations. For combinations involving such operators, two or four competing DNA loop configurations are possible. To unequivocally confirm DNA looping and measure single coherent TALED-driven DNA loops we designed a TALE that recognizes the asymmetrical *lac* O_2_ operator, supporting a single defined binding geometry (Fig. 2A) that can be controlled depending on the orientation of the O_2_ operator. With respect to the operator orientation shown in Fig. 2A, the inverted O_2_ orientation is termed invO_2_ (Fig. 2B).

**Fig. 2.**
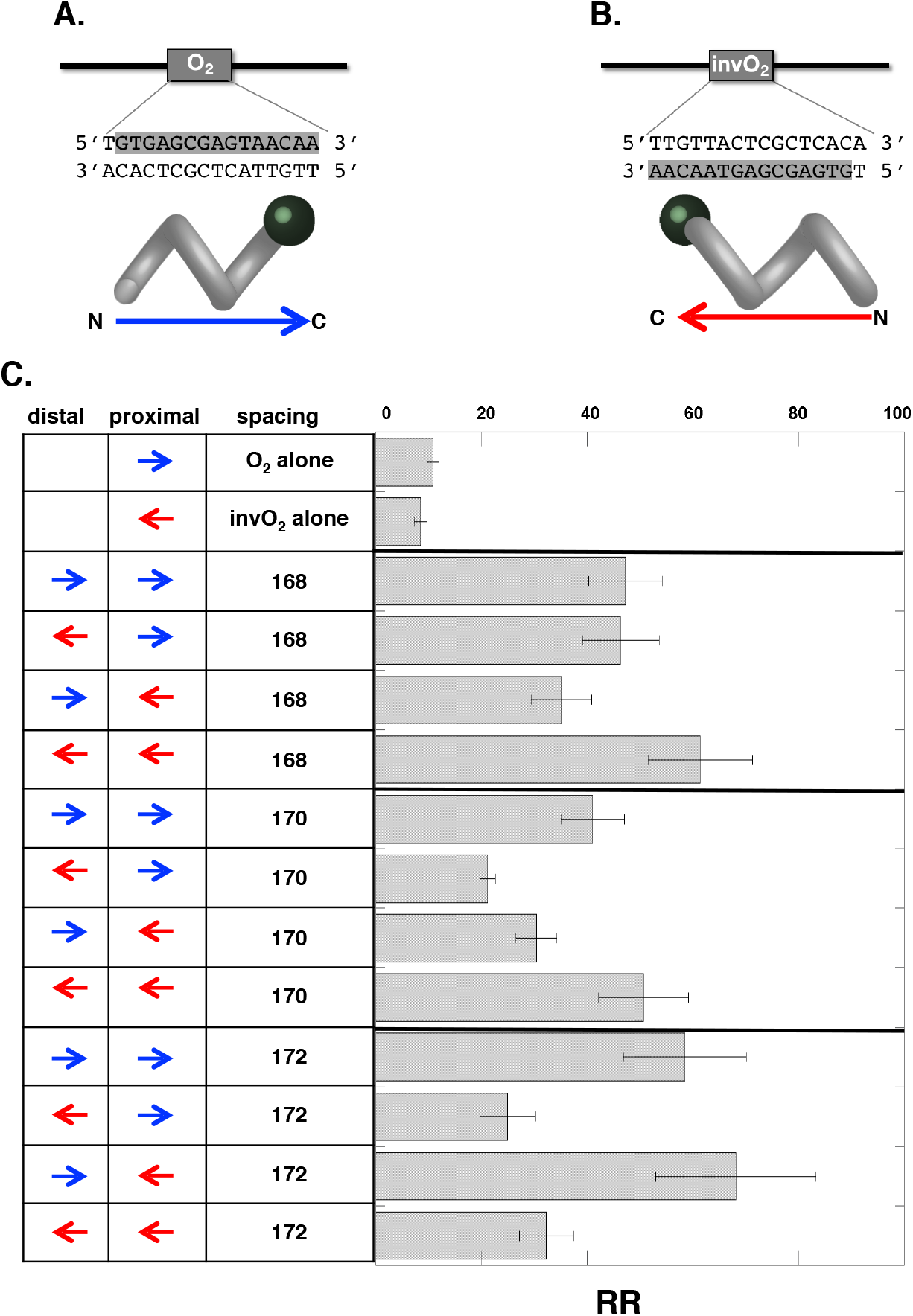
Effect of TALE-operator orientation on repression looping. A. TALE targeting of the purine-rich strand of O_2_ when this asymmetric operator is oriented in the forward direction yields the indicate protein binding polarity. B. The O_2_ operator in a flipped orientation (invO_2_) recognized by the same TALE yields the opposite protein polarity. C. Data comparing repression ratios (as defined in methods) for the indicated operator configurations and center-to-center spacings.

We sought to determine if the stability of a DNA loop driven by the TALED homodimer depends on the relative binding orientations of the two anchoring TALEs (Fig. 2C, left two columns). We therefore collected plasmid-based reporter expression data for three different operator spacings as operator orientations were altered (Supplemental Table S2 and Supplemental Fig. S4). From these results (supplemental Figs. S5), we highlight the repression ratio, *RR*, which compares reporter expression with and without TALED (Fig. 2C). Relative to constructs with a single proximal operator in either orientation, constructs with two operators all showed increased repression (Fig. 2C), consistent with DNA looping (30). Interestingly, loop stability (indicated by extent of reporter repression) varied somewhat as a function of operator orientation, but effects on *RR* were generally less than 2-fold, and there was not a consistent trend that convergent, divergent, or parallel operator orientations were favored. This result suggests that, for the DNA loop sizes studied here, TALED protein flexibility appears to accommodate different loop geometries. In subsequent studies we explicitly distinguish each family of loops (i.e. O_2_-O_2_, invO_2_-O_2_, etc., where the first listed operator is promoter-distal and the second listed operator is promoter-proximal).

### DNA looping by TALE homodimers: effect of operator spacing and context for single loops

Given the apparent similarity of loop stabilities for different operator orientations, we chose to collect a systematic set of new reporter expression data for one representative series of constructs with O_2_-O_2_ and invO_2_-O_2_ operator configurations at different spacings. In the process, we also studied how results were affected by placement of the promoter-reporter construct on the single-copy F’ episome (our conventional choice to mimic the bacterial chromosome) vs. a low copy-number plasmid. Our systematic TALED studies are greatly facilitated in a *ladΓ* background by placement of the promoter-reporter construct on plasmids vs. homologous recombination into the F’ episome. It was therefore important to determine if this more convenient plasmid context gives results comparable to those obtained in the F’ episome. Plasmid-based and episome-based data (Supplemental Tables S3 and S4) measuring repression as a function of operator center-to-center spacing are shown in Fig. 3.

**Fig. 3.**
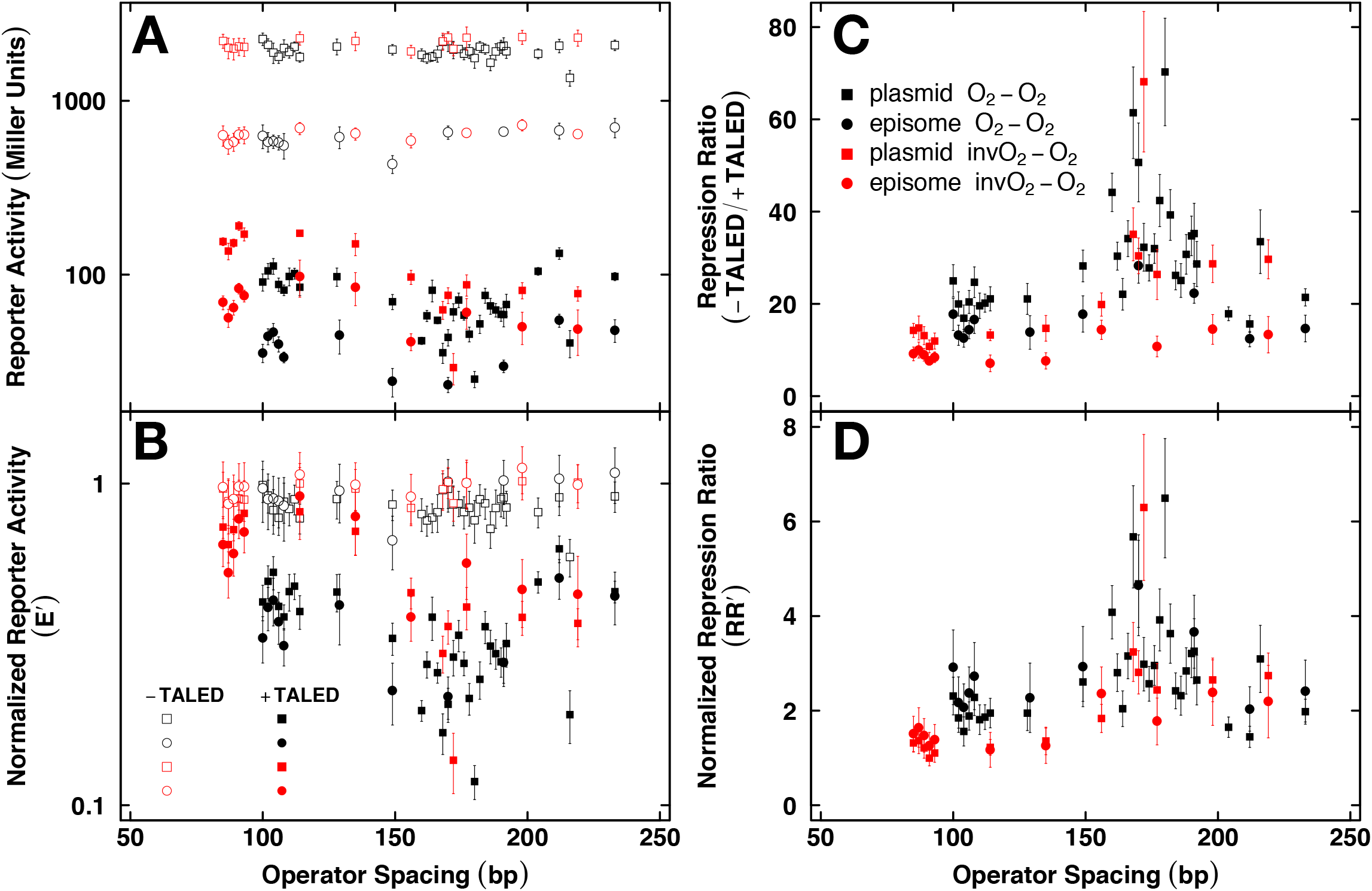
Transcriptional activity and repression by single TALED loop configurations as a function of operator spacing. A. Absolute reporter activity for constructs of the indicated operator pairs at the indicated center-to-center spacings in the indicated context (low-copy plasmid or single-copy episome). B. The data from panel A normalized to control data from corresponding constructs containing only a proximal operator. C. Repression ratio (as defined in methods) for the data in panel A. D. Normalized repression ratio (as defined in methods) summarizing the improvement in repression specifically due to looping.

Panel A of Fig. 3 shows raw reporter activity from the indicated bacterial strains and operator orientations under conditions with or without expression of the homodimer TALED. It is immediately evident that operator pairs lead to length-dependent promoter repression in the presence of TALED protein relative to strains lacking TALEDs (filled vs. open symbols in Fig. 3A). Also evident is the higher reporter activity from cells with plasmid-borne reporters vs. reporters on the single-copy F’ episome (squares vs. circles in Fig. 3A). Both results are consistent with expectations. Interestingly, when reporter activity is normalized to the activity of reference constructs carrying only a single proximal operator (*E’*), the data coalesce into the coherent pattern seen in Fig. 3B. This result indicates that DNA looping behavior is comparable for episomal and plasmid constructs, suggesting that DNA packaging is similar in both contexts, and the titration effect of operator copy number does not substantially influence repression for the intracellular TALED concentration studied here.

TALED-dependent effects are best seen by expressing reporter data as the repression ratio, *RR* (HI 3C), and specific loop-dependent contributions by expressing reporter data as the normalized repression ratio, *RR’* (Fig. 3D), as described in methods. Striking in both panels C and D of Fig. 3 is the evidence for a local maximum in repression in the vicinity of 175-bp operator separation. To analyze and interpret these data more completely, we fit them to an established thermodynamic model of promoter repression by DNA looping [see methods and Supplemental Methods, (10,13,37)]. The results for the O_2_-O_2_ loop series are shown as normalized reporter activity (*E’*) in Fig. 4A and as normalized repression ratio (*RR’*) in Fig. 4B. Results from episome and plasmid contexts are similar enough to be well-characterized by a single set of model parameters (black lines in Fig. 4), clearly demonstrating the characteristic oscillation of repression as a function of operator separation, interpreted as the result of the face-of-the-helix dependence of looping energy favoring repression by untwisted loops. This result firmly establishes DNA looping driven by TALEDs. The comparable, but smaller, invO_2_-O_2_ dataset is similarly analyzed in Fig. 4C and D, with the O_2_-O_2_ model in black for comparison.

**Fig. 4.**
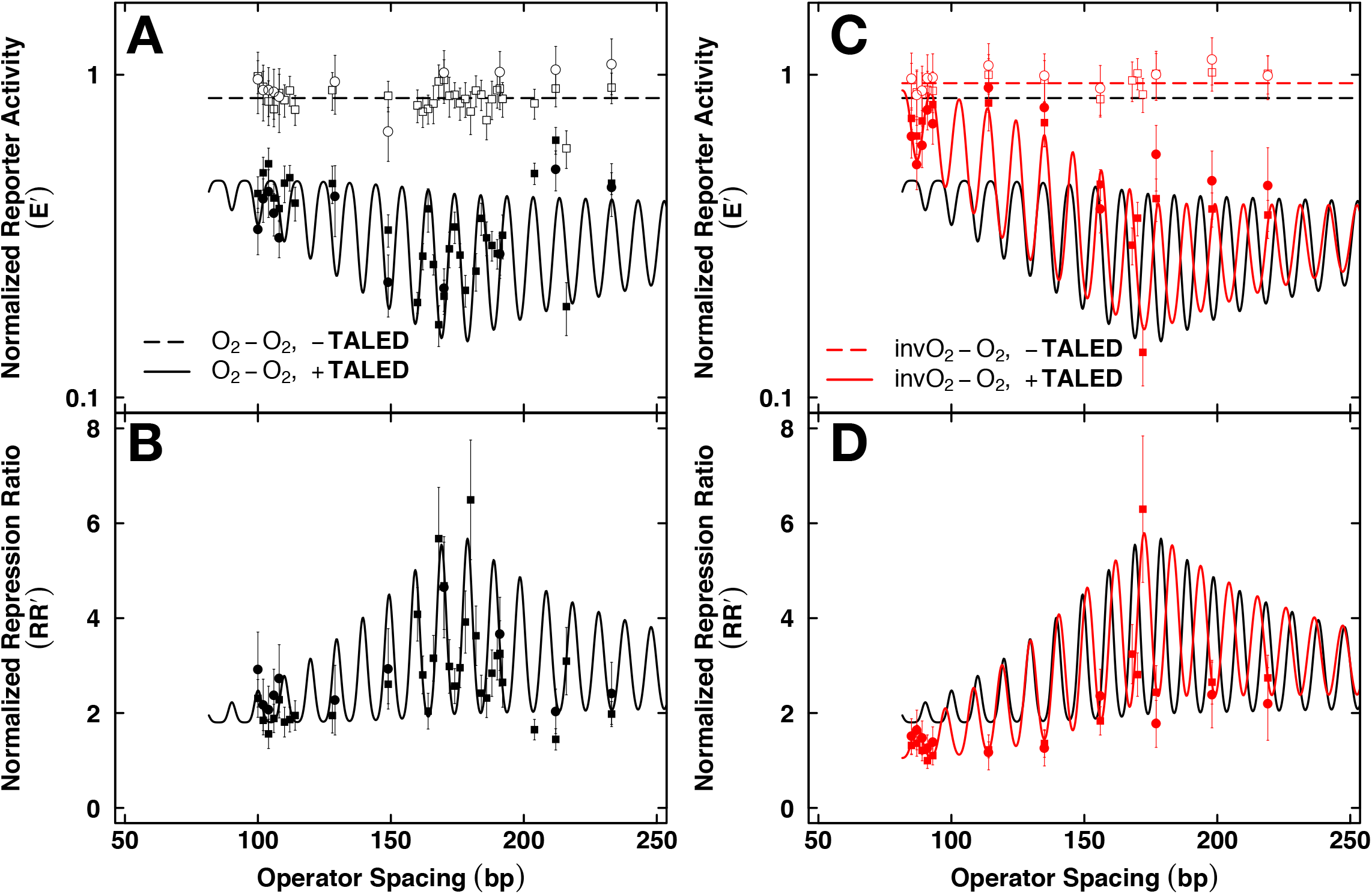
Thermodynamic model fitting as a function of operator spacing for transcriptional activity and repression data from single TALED loop configurations. A. Best fit model for normalized reporter activity data (as defined in methods) for the indicated O_2_-O_2_ operator spacings in the absence or presence of TALED protein. B. Best fit model for normalized repression ratio data of O_2_-O_2_ operator spacings. C. As is panel A except data for invO_2_-O_2_ configurations. Fit from panel A in black for comparison. D. As in panel B but for invO_2_-O_2_ configurations. Fit from panel B in black for comparison.

Values of thermodynamic fitting parameters for these data are shown in Table 1. Several points are worthy of mention. The optimal spacing for repression in each dataset is near 175 bp (~179 and ~172), as expected from visual inspection. Interestingly, for the shortest operator spacings examined, there is greater repression for the O_2_-O_2_ loop series than the invO_2_-O_2_ loop series ^(Fi^g. 4D), which is captured in the model as a greater value of the normalized parameter *K*°_NSL_ (1.24 vs. 0.11, respectively). However, the normalized parameter *K*°_max_ is comparable for both datasets (5.31 vs. 5.72), consistent with the results of Fig. 2 (and Supplemental Table 5) showing that operator orientation has minimal effect on the overall extent of repression. Rather, relative operator orientation appears to have a greater effect on the helical phasing of the loops, with apparent helical repeat changing from 9.86 bp/turn for the O_2_-O_2_ loop series to 10.69 bp/turn for the invO_2_-O_2_ loop series (Fig. 4). The fit value of the torsional modulus of the DNA in the loop (*C*_app_) also captures any additional twist flexibility imparted by the flexible TALED and linker amino acids in the looping process. That estimates of *C*_app_ (1.85 and 0.98) are lower than the common in vitro value of 2.4 (× 10^-19^ erg-cm) (41–43) we attribute to the participation of the flexible TALED protein within the loop, a result that is already well established with LacI looping (10–15). We cannot extract an estimate of the DNA persistence length *per se* from the model without knowing the extent of DNA bending in the loop (see Supplemental Methods). However, the empirical fit parameter *P_app_* [which includes contributions from DNA persistence length (*P*), the thermal energy (RT), and the extent of bending (ΔΘ) in a single constant] captures the variable level of repression as a function of operator spacing. In the case of maximal repression, this corresponds to a bending energy of the loop of 2.4 RT and 1.9 RT, respectively for the two datasets. A similar value has been observed for LacI repression looping (2.3 RT), despite maximal repression occurring from a smaller operator spacing.

**Table 1.** Thermodynamic model fits for data with a single loop configuration

|  | TALED<sup>a</sup> |  |  | WT LacI<sup>b</sup> |  |
| --- | --- | --- | --- | --- | --- |
| Parameter | O<sub>2</sub>-O<sub>2</sub> | invO<sub>2</sub>-O<sub>2</sub> |  | Osym-O2, -IPTG | Osym-O2, +IPTG |
| $hr$ (bp/turn) | $9.86 \pm 0.11$ | $10.69 \pm 0.11$ | | $11.44 \pm 0.74$ | $10.73 \pm 0.49$ |
| $C_{app}$ ($\times 10^{-19}$ erg-cm) | $1.85 \pm 0.33$ | $0.98 \pm 0.21$ | | $0.76 \pm 0.42$ | $0.64 \pm 1.11$ |
| $K_{max}^o$ | $5.31 \pm 0.69$ | $5.72 \pm 1.37$ | | 68.62 | 17.85 |
| $K_{NSL}^o$ | $1.24 \pm 0.24$ | $0.11 \pm 0.18$ | | 10.45 | 0 |
| $sp_{optimal}$ (bp) | $178.87 \pm 0.32$ | $172.41 \pm 0.50$ | | $78.27 \pm 0.53$ | $78.82 \pm 0.46$ |
| $P_{app}$ (bp) | $420.57 \pm 73.81$ | $337.33 \pm 68.10$ | | $15.71 \pm 150.54$ | $184.66 \pm 185.59$ |
|  | episome | episome |  | episome | episome |
| $K_O$ | 5.56 | 5.78 | | 2.45 | 0.13 |
| $K_{max}$ | 29.49 | 33.02 | | $167.78 \pm 60.60$ | $2.39 \pm 0.54$ |
| $K_{NSL}$ | 6.91 | 0.65 | | $25.55 \pm 31.29$ | $0.00 \pm 2.06$ |
| $K_{O2}$ [-TALED] | 0.08 | 0.12 | | | |
| $K_{NS}$ [-TALED] | 0.08 | 0.02 | | | |
|  | plasmid | plasmid |  |  |  |
| $K_O$ | 9.82 | 10.01 | | | |
| $K_{max}$ | 52.14 | 57.19 | | | |
| $K_{NSL}$ | 12.21 | 1.13 | | | |
| $K_{O2}$ [-TALED] | 0 | 0.02 | | | |
| $K_{NS}$ [-TALED] | 0.18 | 0.06 | | | |

<sup>a</sup>All parameters are +TALED except when otherwise indicated.
<sup>b</sup>The normalized parameters were not used for fitting in Becker et al. 2005 (10).

Any repression looping system based on dimeric proteins that simultaneously bind two DNA sites will show a repression optimum that depends on the concentration of the protein dimer. This occurs because, at high concentrations, preformed dimers may saturate both DNA sites without looping. Future experiments to estimate and systematically alter in vivo TALED concentrations will allow exploration of this variable.

### Global analysis of TALED and LacI repression loops

Because of our past experience analyzing repression loops by LacI repressor (10–15), mixed competing TALED loop configurations (30), and the single TALED loop configurations described in this work, we have the opportunity for global comparison of the repression loop properties based on our data for these systems. Examples from each of these datasets are shown together in Fig. 5A and B. The Supplemental Methods (and Supplemental Figs. S6 – S8) describe how model predictions constructed using the O_2_-O_2_, invO_2_-O_2_, O_2_-invO_2_, and invO_2_-invO_2_ datasets without adjusting any parameters yield results that qualitatively match the previous O_sym_-O_sym_, O_sym_-O_B_, and O_sym_-invO_B_ datasets, despite differences in the identity of the TALED, DNA-binding affinities, operator spacings, and whether the reporter was plasmid-borne or episome-borne.

**Fig. 5.**
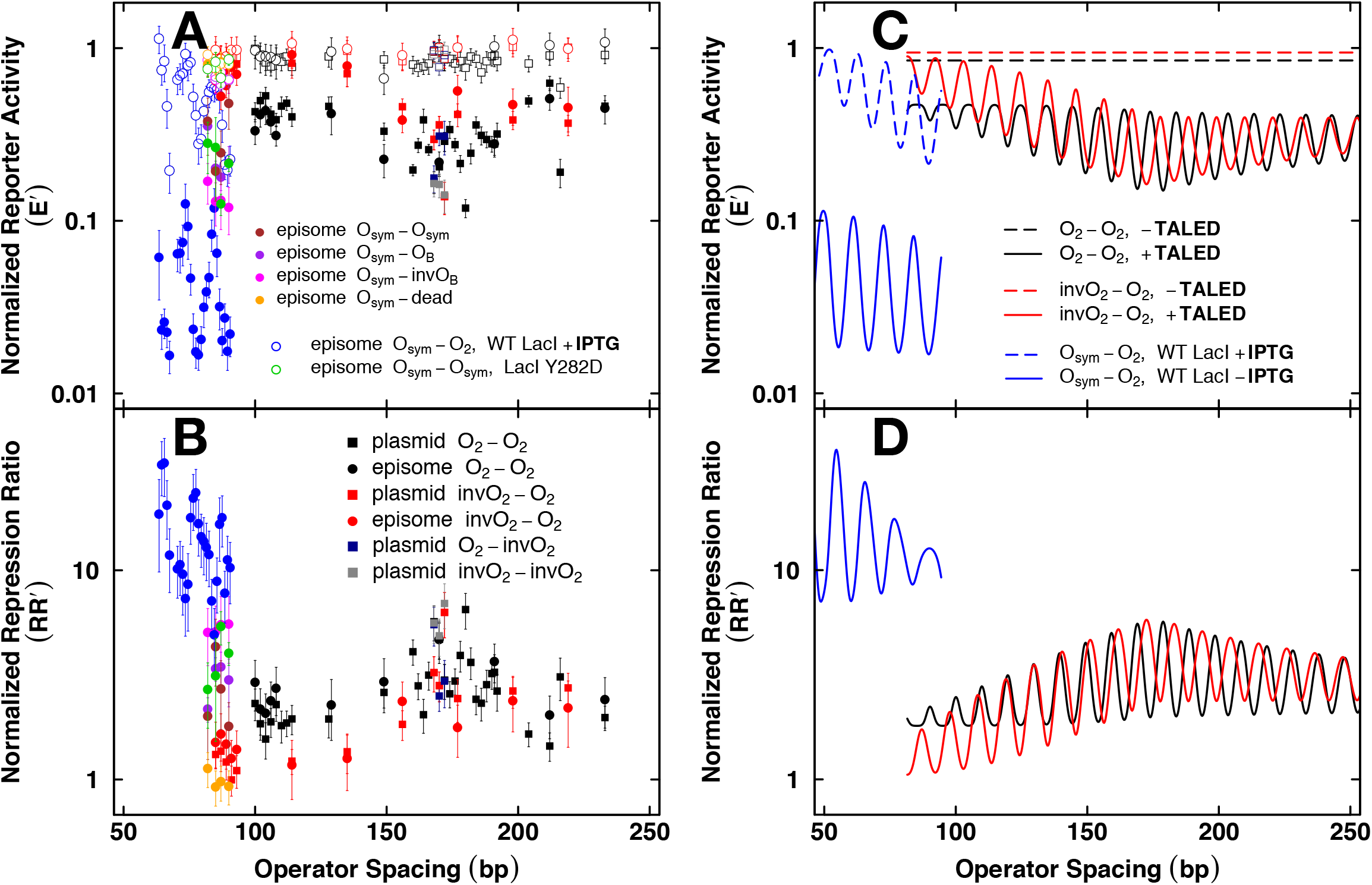
Compilation of repression data for DNA loops anchored by TALEDs or LacI repressor and comparison of thermodynamic model fits for data from single configuration *lac* and TALED loops. A. Normalized reporter activity for all indicated constructs from this and prior published reports from our laboratory, including LacI repressor loops and all TALED single loop configurations (involving only O_2_ and invO_2_ operators) and multiple competing TALED loop configurations (involving O_sym_). B. Data plotted as normalized repression ratio. C. Best fit model for reporter activity from the indicated cases. Black indicates fits for O_2_-O_2_ operator spacings with TALED. Red indicates fits for invO_2_-O_2_ operator spacings with TALED. Blue indicates fits for Osym-O_2_ operator spacings with LacI repressor. D. Model fits for normalized repression ratio.

Comparing to our previous *lac* studies, a direct example is provided by episome-based O_sym_-O_sym_ constructs. The brown circles in Fig. 5A and B show reporter activity for four operator spacings near 86 bp with or without TALED protein (filled vs. open symbols). The green circles in Fig. 5A and B show reporter activity from an *E. coli* strain with wild-type lac repressor (WT LacI, filled symbols) or a strain with a totally disabled lac repressor (LacI Y282D, open symbols). The latter is equivalent to the absence of TALED so that the maximal promoter activity in the absence of functional protein is the same for both. The extent of decreased report activity caused by protein-mediated looping (WT LacI, green vs. TALED, brown) is very similar (filled symbols in Fig. 5A), despite different anchoring proteins. Even though this particular operator configuration leads to very high levels of total repression (several hundred-fold), the contribution from DNA looping is only modest (*RR’* value of 2-6 in Fig. 5B) since each protein binds the operator tightly even in the absence of a distal operator (*RR* of 113 vs. 75, respectively for WT LacI vs. TALED).

To better observe the effects of looping on repression, a majority of the previous work with lac repressor explored the combination of a strong distal operator sequence (Osym) with a weak proximal operator sequence (O_2_). In an episome, the proximal O_2_ alone with wild type LacI accounted for an *RR* value of ~3 relative to the absence of repressor (i.e. LacI Y282D mutant). A similar value was observed for WT LacI assayed with or without IPTG inducer. *RR* values for proximal O_2_ alone with TALED are slightly larger, ~6 (episome) vs. ~11 (plasmid), and vary less then 2-fold based on reporter type. The blue circles (open vs. filled) in Fig. 5 display lac repressor data (±IPTG inducer) for episomal constructs with operator spacings ranging from 60 to 90 bp. Here several important observations can be made. From a total repression enhancement up to 100-fold, lac repressor-mediated looping is the dominant contributor with an *RR’* of 5-33 and a mean of ~15 (Fig. 5B), in contrast to the maximal TALED *RR’* of ~6. For clarity, Fig. 5C and D display the thermodynamic model fits to the different datasets with a single loop configuration. By comparing the fit value of the normalized parameter *K*°_max_ (Table 1), it is evident that the contribution of looping to repression is higher for lac repressor (~69) than for TALED (~6). However, each of the distal operator sequences was weak in the TALED datasets, with *RR* of either ~11 (proximal O_2_ alone, plasmid) or ~8 (proximal invO_2_ alone, plasmid). Future creation of heterodimer TALEDs capable of binding both strong distal and weak proximal operator sequences will further tease out this effect.

### Comparing repression by lac repressor and by TALEDs

We set out to apply artificial TALED proteins in the context of a set of promoter-reporter constructs previously assembled to study DNA looping by LacI repressor in vivo. While this approach created some complications in our initial report (binding of the initial TALEs to the symmetrical *lac* O_sym_ operator can occur in either of two orientations), we now have designed TALEs that recognize asymmetrical operators so that single defined loops are created using distinct orientations of the O_2_ operator sequence. This has allowed detailed measurement of the defined TALED loop system in living bacteria, facilitating comparison with loops driven by LacI repressor. As shown in ^Fi^g. 5C and D, the classic oscillation pattern in our new data make it unequivocal that the TALED system drives DNA looping. Fig. 5D summarizes the observation that the contribution of looping to repression is higher for lac repressor than for TALEDs. The two systems have similarities and differences.

With respect to similarities, loops driven by LacI or TALEDs have in common that the optimal loop length is far smaller than expected for naked DNA (near four persistence lengths). Over the length scales we previously studied for LacI, DNA bending appears to be a much smaller obstacle than expected for looping, with DNA twist energy playing a more obvious role (10). We have interpreted this as evidence for the effect of architectural DNA binding proteins in vivo (10–12) or location of the repression loop at the apex of a supercoiled plectonemic domain.

With respect to differences, our datasets show strong LacI repressor looping even for loops smaller than 100-bp in length, in agreement with prior studies (20), whereas TALED looping is optimal closer to 175 bp. Several considerations may explain the different behavior of the two systems. First, whereas the LacI tetramer is a stable dimer of dimers, TALE dimerization via the FKBP(F36M) domain is weak, with an equilibrium dissociation constant in the tens of micromolar. This implies that loops involving expensive DNA deformation may not be supported by the weak TALE dimerization interface. This concept would explain relatively low repression and little torsional dependence for short operator spacings, then a regime of increasing repression that depends on the second operator with gradually decreasing torsional oscillation as operator spacing increases. Second, the concept of a small molecule chemical inducer is different between the systems. Whereas *lac* induction involves a small metabolite that decreases repressor affinity for its DNA operators, TALED anti-repression by small molecules in our engineered system does not change operator affinity, but alters protein dimerization (30). Third, occupancy of the promoter-proximal operator by a TALE protein, even if TALE dimerization has been blocked, results in higher basal repression for TALE monomer binding to this operator than for the LacI weakened by inducer binding. Fourth, although LacI binding undoubtedly changes the physical properties of the occupied operator DNA (44–46), DNA recognition by TALE protein wrapping of the operator major groove is expected to confer additional rigidity (32), constraining operator conformations within repression loops. This consideration, together with weak TALE dimerization, may explain the large optimal loop lengths observed in the TALED system. Thus, whereas the apparent optimal DNA loop length for lac repressor was found to be near 80 bp (10,20), the maximum for TALEDs appears to be near 175 bp (Fig. 5).

The concept of an optimal DNA length for protein-mediated looping is, in itself, interesting. It is possible that looping optima reflect the same principles embodied in predictions of the Wormlike Chain polymer model for DNA cyclization (17,47). Because of its high relative stiffness, the effective end-end concentration *(J)* is extremely low for short DNA, rising rapidly to an optimal cyclization DNA length, before gradually falling for longer lengths because of entropic effects. We suggest that these principles are also revealed for loops mediated by LacI repressor or TALEDs. Our modeling of TALED looping therefore combines the two effects to create a looping probability maximum such that for spacings shorter than 175 bp an enthalpy penalty limits looping and for spacings larger than 175 bp an entropy penalty reduces looping. While successfully fitting the data, the physical interpretation of the resulting empirical parameter *P*_app_ is now more difficult. Without knowing the actual extent of DNA bending in the loop it is not possible to extract a conventional value of *P*. Considerations based on the Wormlike Chain polymer model (48) allow estimation of apparent values of the DNA persistence length (which contains contributions from both proteins and negatively supercoiled DNA in vivo). These estimated persistence lengths are ~11 nm for LacI repressor and ~17 nm for the TALE homodimer studied here, contrasting with ~50 nm for DNA in vitro. The different optimal looping DNA lengths may reflect the considerations raised above, or even the possibility that protein-DNA loops have evolved to recruit architectural DNA binding proteins that reduce apparent DNA stiffness (10–12,16).

## Conclusion

The current work is the first to create unique DNA repression loops using programmable Transcription Activator-Like Effector dimer (TALED) proteins in *E. coli.* Systematic study of repression as a function of loop size, with data fitting to a thermodynamic model, now allows this system to be compared in detail with lac repressor loops, and relevant biophysical parameters to be estimated. This approach has implications for the artificial regulation of gene expression.

## Author contributions

Project conception and manuscript preparation: NAB, JPP, KJC, LJM

Experimental work: NAB, JPP, TLS, JPW, WJP

## Acknowledgments

We thank members of the Maher laboratory for assistance.

